# Integrated single-nucleus and spatial transcriptomics captures transitional states in soybean nodule symbiosis establishment

**DOI:** 10.1101/2022.06.30.498286

**Authors:** Zhijian Liu, Xiangying Kong, Yanping Long, Hong Zhang, Jinbu Jia, Lijuan Qiu, Jixian Zhai, Zhe Yan

## Abstract

Legumes form symbiosis with rhizobium leading to the development of nitrogen-fixing nodules. By integrating single-nucleus and spatial transcriptomics, we established a cell atlas of soybean nodules and roots. In central infected zone of nodule, we found that uninfected cells specialize into functionally distinct sub-groups during nodule development and revealed a transitional subtype of infected cells with enriched nodulation-related genes. Overall, our results provide a single-cell perspective for understanding rhizobium-legume symbiosis.

## Main

On compatible host plants, the rhizobium bacteria infect and form symbiotic organ-nodules in the root, establishing nitrogen-fixing nodules which can convert atmospheric nitrogen into organic ammonia for host plant development. Despite remarkable progress by molecular genetics have established the framework of the nodulation and symbiotic nitrogen fixation (SNF)^1^, our understanding of cellular heterogeneity and developmental lineage of nodule is still limited.

To reveal the cell-type specific dynamic gene expression during nodule development in soybean, we established three single-nucleus libraries with two different developmental stages of nodules (at 12 days post-infection (dpi) and 21 dpi) and the corresponding region of roots where the nodules were formed at 21 dpi as control (Fig.1a). We obtained a total of 26712 high quality single-nucleus transcriptomes in the three libraries, which covering 39337 genes, with median genes/nucleus at 1342 and median UMIs/nucleus at 1636 (Supplementary Data 1). After integration of the three datasets using scVI^2^, we obtained 15 cell clusters (Fig.1b-c) and a series of up-regulated genes for each cluster (Supplementary Fig.1, Supplementary Data 2). With known soybean marker genes, orthologs of marker genes in Arabidopsis as well as public Arabidopsis scRNA-seq dataset^3^, we successfully identified root epidermis (cluster 5), root vascular bundle (cluster 3), nodule vascular bundle (cluster 9), nodule cortex (cluster 1) and infected cells (ICs) in nodule central infected zone (CIZ) (cluster 12) (Supplementary Fig.2-4, Supplementary Data 3). However, due to the scarcity of marker genes in soybean nodule, there are still many cell clusters cannot be successfully assigned, especially those dominated by nodules (Supplementary Fig.4b). To overcome this problem, we used stereo-seq^4^ to track the spatial expression of genes of the same developmental stage nodules (Fig.1a, Supplementary Fig.5). By performing a deconvolution-based approach on these spatial transcriptomes, we validated the cluster identities which we detected above and further assigned cluster 0 (in CIZ), 2 (outer cortex), 4 (outer cortex), 7 (in CIZ) and 11 (in CIZ) based on their distribution over space (Fig.1c-d). To validate our final annotation, we performed GUS staining and RNA *in situ* hybridization with four genes specific expressed in CIZ, vascular bundle and inner cortex, and observed corresponding cell-type specific signals in nodules (Fig.1e, Supplementary Fig.6). In summary, here we successfully classified the major cell types of both root and nodules.

**Figure 1.**
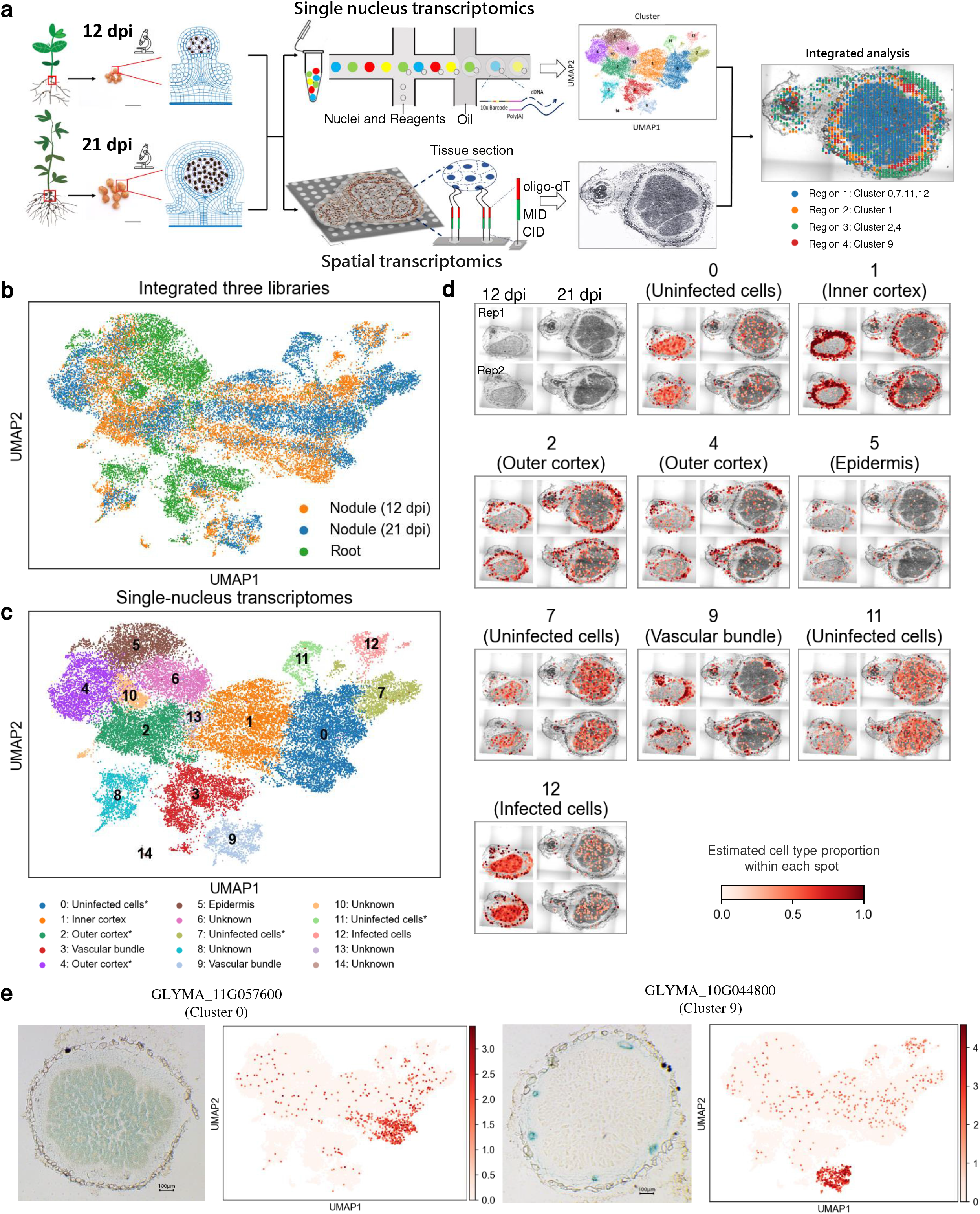
Combined spatial transcriptomes and single-nucleus transcriptomes reveal nodule heterogeneity at different developmental stages. **a**. Schematic diagram of the integration of single-nucleus and spatial transcriptomics analysis. **b**. Integration of three single-nucleus datasets. **c**. UMAP visualization of identified 15 cell clusters in nodules and roots. “*” indicates that the cluster is annotated by spatial transcriptome. **d**. Spatial distribution of different cell types in the bright field picture. Upper left, bright field image of nodule sections used to prepare the spatial transcriptome. Two replicates are analyzed for both 12-dpi and 21-dpi nodules. Others, spatial distribution of cell type proportions for each single-nucleus cluster. The colors represent the fraction of single-nucleus transcriptomes of each cluster deconvolved by destVI. **e**. Validation of annotation results by GUS-reporter lines.

There are four clusters 0, 7, 11 and 12 co-localized in CIZ of nodule (Fig.1d). By examining the expression of orthologs which are up-regulated in *Lotus japonocus* uninfected cells (UCs) and infected cells (ICs) ^5^, we identified cluster 0, 7, 11 as UC and further confirmed cluster 12 as ICs (Fig.2a, Supplementary Fig.7, Supplementary Data 4). In UCs, cluster 0 is shared by nodules at two different developmental stages, while two clusters (7, 11) are almost only found in 21-dpi nodule cells (Fig.2b). To reveal the differentiation trajectory of UC cells, we performed pseudo-time analysis and found that cluster 7 and 11 are developed from cluster 0, indicating differentiation of functions of UC cells during maturation (Fig.2c). In tropical legumes like soybean, ureides are the primary export forms in root nodules from currently fixed nitrogen. It was reported that ureides are mainly synthesized in UCs and enzymes which are responsible for ureides biosynthesis present a higher specific activity in the UCs^6,7^. For ureide biogenesis, the uricase and aspartate aminotransferase genes which expressed in nodules are expressed in all three UC clusters and especially up-regulated in UC cluster 7 (Fig.2d). While for ureide transportation, 2 of 3 ureide permease genes are mainly expressed in UC cluster 0 (Fig.2d). These results revel a complex compartmentalization in UCs during ureide production and transportation in soybean nodules. Moreover, we found that expression of six of eight beta amylase genes is significantly up-regulated in cluster 11 (Fig.2d) and the pathways associated with polysaccharide catabolic process, starch catabolism are also activated, which indicated that cluster 7 involved in energy supply for symbiotic nitrogen fixation (Supplementary Fig.8). Taken together, these results revealed that the UCs continue to differentiate into functionally specialized sub-cell types during development, which can facilitate the exchange of nutrient and energy sources required for symbiosis.

**Figure 2.**
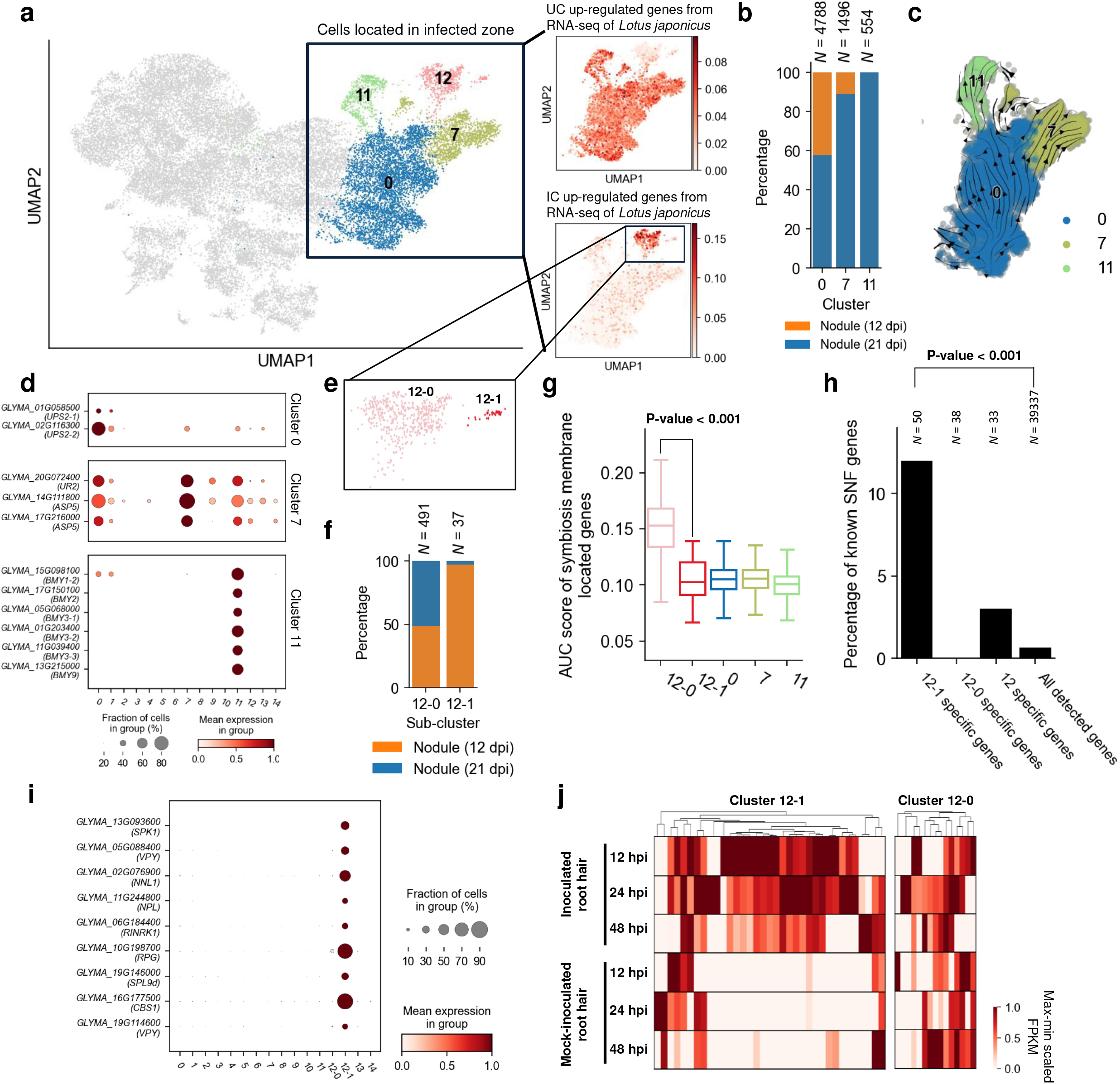
Dissection central infected zone reveals distinct subtypes of nodule cells. **a**. The distribution of orthologs of UCs and ICs highly expressed genes in *Lotus japonicus* in the UMAP. Expression levels of gene sets are measured by AUC score. **b**. Bar chart representing the percentage of cells from different samples in each UC clusters. N indicates the cell number. **c**. Developmental trajectories of UCs inferred using cellrank and cytotrace. Colors represent different IC clusters (0, 7, 11). **d**. Dotplot representing the expression pattern of representative up-regulated genes for each UC cluster. **e**. UMAP visualization of identified IC subclusters. **f**. Bar chart representing the percentage of cells from different samples in each sub-cell type. N indicates the cell number. **g**. AUC score of genes encoding symbiosis membrane protein. The p-values are calculated by the Mood’s median test. The p-values between 12-1 and the remaining three UC clusters are greater than 0.05. **h**. Percentage of known symbiotic nitrogen fixation genes collected by Roy *et al*. in different cell-type-specific gene sets. The calculation of the P-value is detailed in Supplementary Fig.11. N indicates the gene number. **i**. Dotplot representing the expression pattern of 12-1-specific known symbiotic nitrogen fixation genes. **j**. Heatmap representing expression pattern of detected specific genes for subcluster 12-0 and 12-1 in inoculated and mock-inoculated root hair datasets.

We then focused on infected cells, the core sites of SNF. Consistent with previous reports, some reported IC-specific genes, such as *GmSYMREM*^8^, *GmN56*^9^, *GmENOD55*^10^ are all restricted in cluster 12 and leghemoglobin genes^11^ are up-regulated in cluster 12 (Supplementary Fig.9). The gene ontology (GO) analysis of up-regulated genes in ICs showed that the pathways associated with carbohydrate transmembrane transport and nitrogen-containing amino acid synthesis are activated, presenting the active carbon and nitrogen exchange between soybean and rhizobia in ICs (Supplementary Fig.10). By re-clustering of ICs, we found that they could be further divided into two sub-cell type (subcluster 12-0 and 12-1) (Fig.2e). Subcluster 12-0 is shared by nodules at two different developmental stages but the small subcluster 12-1 is almost exclusively occupied by the 12-dpi immature nodule (Fig.2f). The expression levels of genes encoding symbiosome membrane protein^12^ were much higher in cluster 12-0 than in subcluster 12-1, while there was almost no difference between subcluster 12-1 and UCs (Fig.2g), indicating a more active movement of solutes between symbionts in subcluster 12-0 and subcluster 12-1 is a transitional cell type of ICs during nodule development. We checked subcluster 12-1 specific gene and found nearly 12% of the genes (6/50) are included in known SNF genes collected by Roy *et al*^1^. This proportion is significantly higher than cluster 12, 12-0 and all detected clusters (Fig.2h, Supplemental Fig.11). Besides, 3 of the remaining 44 genes are reported as SNF genes recent years^13,14^ (Fig.2h-i, Supplementary Data 2). We further found that all these 9 SNF genes, including *SPK1*^15^, VPY^16^, *NNL1*^*13*^, *NPL*^17^, *RINRK1*^18^, *RPG*^19^, *SPL9d*^14^, *CBS1*^20^, are involved in the formation of infection threads (ITs). ITs take place in root hair after rhizobium attachment and they assist rhizobium reach and finally release into developing nodules. We analyzed the expression of 12-1 cluster genes in soybean root hair in the early stage of rhizobial infection (12-dpi, 24-dpi and 48-dpi) and 60% of these genes (21/35) are expressed only after rhizobia inoculation^21^(Fig.2j). In contrast, of the cluster 12-0 specific genes, only 2 were induced after induction. These results imply that cluster 12-1 could involve in ITs extension and the rhizobia release in ICs during nodule maturation and genes that are specifically expressed in 12-1 may play a critical role in interaction between soybean and rhizobium in different stages of symbiosis establishment.

Overall, we provide a comprehensive cellular atlas by combining single-cell data with spatial transcriptomic data. Based on this atlas, we identified rare cell subtypes and revealed their distinct roles for nodule maturation and function. To help community to explore the heterogeneity of different cell types in soybean nodules, we also present a web server (http://159.138.151.218:3569/) to facilitate the use of the datasets generated in this study. In conclusion, we provide a data resource that will contribute to learning the regulatory network of nodule development at the single cell level in the future.

## Materials and Methods

Please refer to Supplementary material.

## Data Availability

The raw sequencing data generated in this study were deposited in Chine National Center for Bioinformation with accession PRJCA009893 (reviewer link: https://ngdc.cncb.ac.cn/gsa/s/1rNqwyk1).

## Acknowledgments

The group of Z.Y. is supported by the Strategic Priority Research Program of the Chinese Academy of Sciences, Grant No. XDA24010205 and XDA28030101; Agricultural Science and Technology Innovation Program. The group of J.Z. is supported by the National Key R&D Program of China Grant (2019YFA0903903); an NSFC to J.Z. (31871234); the Shenzhen Sci-Tech Fund (KYTDPT20181011104005); the Key Laboratory of Molecular Design for Plant Cell Factory of Guangdong Higher Education Institutes (2019KSYS006) and the Stable Support Plan Program of Shenzhen Natural Science Fund Grant (20200925153345004).

## Author Contributions

Z.Y. and J.Z. designed the experiments. Z.L., X.K. and Y.L. performed the experiments. Z.L. and Y.L. analyzed the data. Z.Y., J.Z., Z.L., X.K. and Y.L. wrote the manuscript, L.Q. provided conceptual insight. H.J. and J.J. edited the article.

## Competing interests

The authors declare no competing interests.

## Notes

### Competing Interest Statement

The authors have declared no competing interest.

## References

1 Roy, S. et al. Celebrating 20 years of genetic discoveries in legume nodulation and symbiotic nitrogen fixation. The Plant Cell 32, 15–41 (2020).

2 Lopez, R., Regier, J., Cole, M. B., Jordan, M. I. & Yosef, N. Deep generative modeling for single-cell transcriptomics. Nature Methods 15, 1053–1058 (2018).

3 Shahan, R. et al. A single-cell Arabidopsis root atlas reveals developmental trajectories in wild-type and cell identity mutants. Developmental Cell 57, 543–560. e549 (2022).

4 Xia, K. et al. The single-cell stereo-seq reveals region-specific cell subtypes and transcriptome profiling in Arabidopsis leaves. Developmental Cell 57, 1299–1310. e1294 (2022).

5 Wang, L. et al. Single cell-type transcriptome profiling reveals genes that promote nitrogen fixation in the infected and uninfected cells of legume nodules. Plant Biotechnology Journal 20, 616–618 (2022).

6 Newcomb, E. H. & Tandon, S. R. Uninfected cells of soybean root nodules: ultrastructure suggests key role in ureide production. Science 212, 1394–1396 (1981).

7 Hanks, J. F., Schubert, K. & Tolbert, N. Isolation and characterization of infected and uninfected cells from soybean nodules: role of uninfected cells in ureide synthesis. Plant Physiology 71, 869–873 (1983).

8 Fan, W. et al. Rhizobial infection of 4C cells triggers their endoreduplication during symbiotic nodule development in soybean. New Phytologist 234, 1018–1030 (2022).

9 Kouchi, H. & Hata, S. GmN56, a novel nodule-specific cDNA from soybean root nodules encodes a protein homologous to isopropylmalate synthase and homocitrate synthase. Molecular Plant-microbe Interactions: MPMI 8, 172–176 (1995).

10 de Blank, C. et al. Characterization of the soybean early nodulin cDNA clone GmENOD55. Plant Molecular Biology 22, 1167–1171 (1993).

11 Appleby, C. A. Leghemoglobin and Rhizobium respiration. Annual Review of Plant Physiology 35, 443–478 (1984).

12 Luo, Y., Liu, W., Sun, J., Zhang, Z.-R. & Yang, W.-C. Quantitative proteomics reveals key pathways in the symbiotic interface and the likely extracellular property of soybean symbiosome. Journal of Genetics and Genomics (2022).

13 Zhang, B. et al. Glycine max NNL1 restricts symbiotic compatibility with widely distributed bradyrhizobia via root hair infection. Nature Plants 7, 73–86 (2021).

14 Yun, J. et al. The miR156b-GmSPL9d module modulates nodulation by targeting multiple core nodulation genes in soybean. New Phytologist 233, 1881–1899 (2022).

15 Liu, J., Liu, M. X., Qiu, L. P. & Xie, F. SPIKE1 activates the GTPase ROP6 to guide the polarized growth of infection threads in Lotus japonicus. The Plant Cell 32, 3774–3791 (2020).

16 Murray, J. D. et al. Vapyrin, a gene essential for intracellular progression of arbuscular mycorrhizal symbiosis, is also essential for infection by rhizobia in the nodule symbiosis of Medicago truncatula. The Plant Journal 65, 244–252 (2011).

17 Xie, F. et al. Legume pectate lyase required for root infection by rhizobia. Proceedings of the National Academy of Sciences 109, 633–638 (2012).

18 Li, X. et al. Atypical receptor kinase RINRK1 required for rhizobial infection but not nodule development in Lotus japonicus. Plant Physiology 181, 804–816 (2019).

19 Arrighi, J.-F. et al. The RPG gene of Medicago truncatula controls Rhizobium-directed polar growth during infection. Proceedings of the National Academy of Sciences 105, 9817–9822 (2008).

20 Sinharoy, S. et al. A Medicago truncatula cystathionine-β-synthase-like domain-containing protein is required for rhizobial infection and symbiotic nitrogen fixation. Plant Physiology 170, 2204–2217 (2016).

21 Libault, M. et al. Complete transcriptome of the soybean root hair cell, a single-cell model, and its alteration in response to Bradyrhizobium japonicum infection. Plant Physiology 152, 541–552 (2010).

