## supplemental Figure for "Integrated single-nucleus and spatial transcriptomics captures transitional states in soybean nodule symbiosis establishment"

### Slide 1
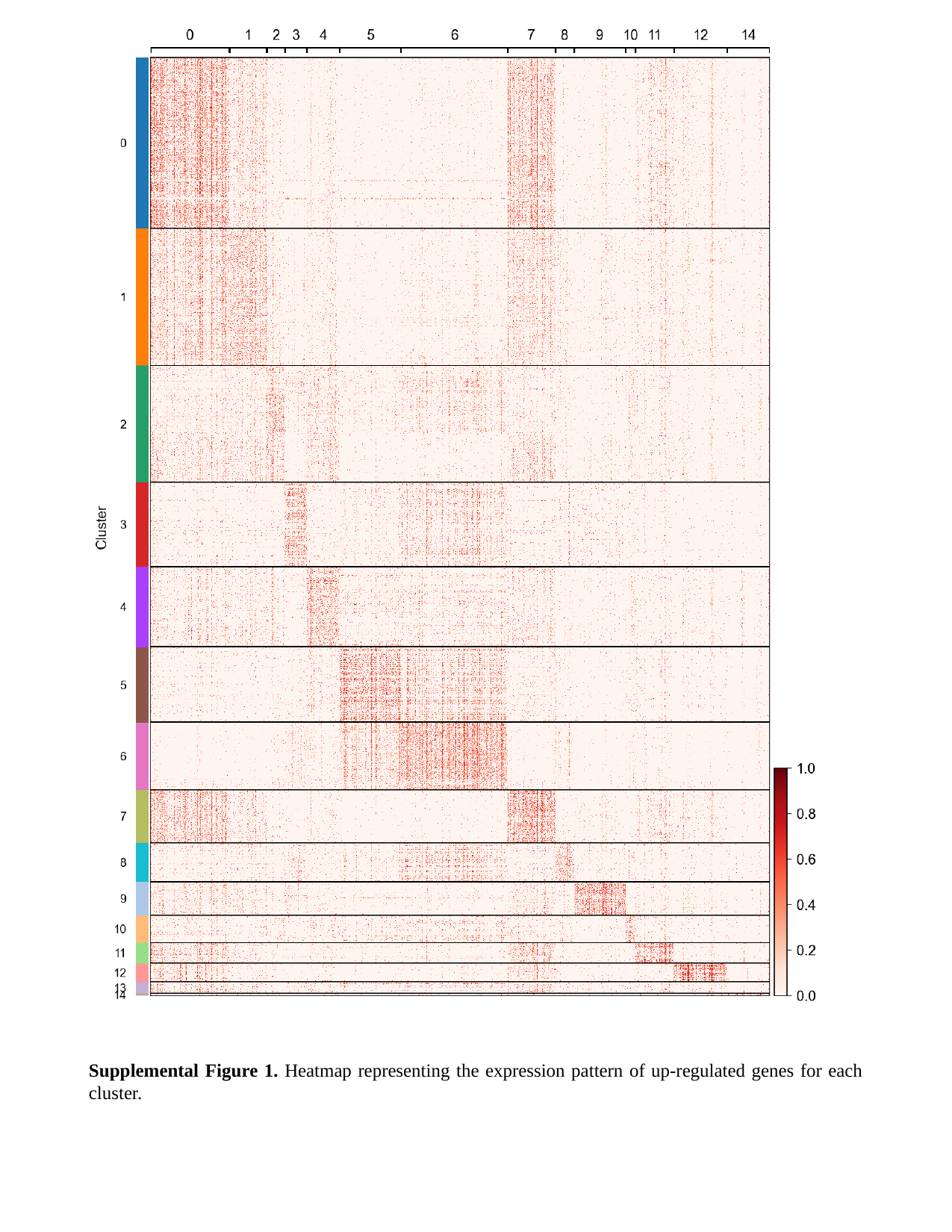

Supplemental Figure 1. Heatmap representing the expression pattern of up-regulated genes for each cluster.

### Slide 2
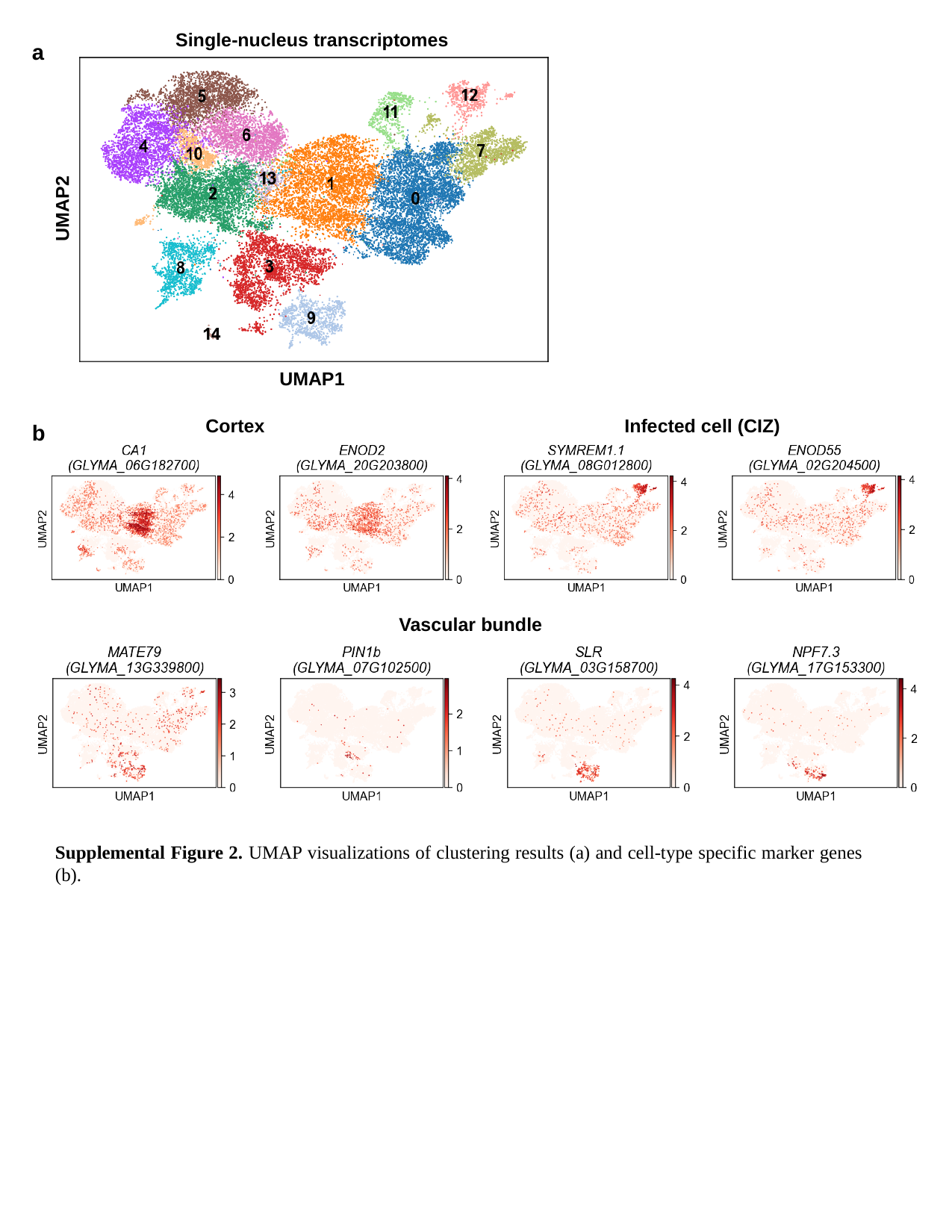

Single-nucleus transcriptomes
UMAP2
UMAP1
a
Cortex
Infected cell (CIZ)
b
cortex
Vascular bundle
Supplemental Figure 2. UMAP visualizations of clustering results (a) and cell-type specific marker genes (b).

### Slide 3
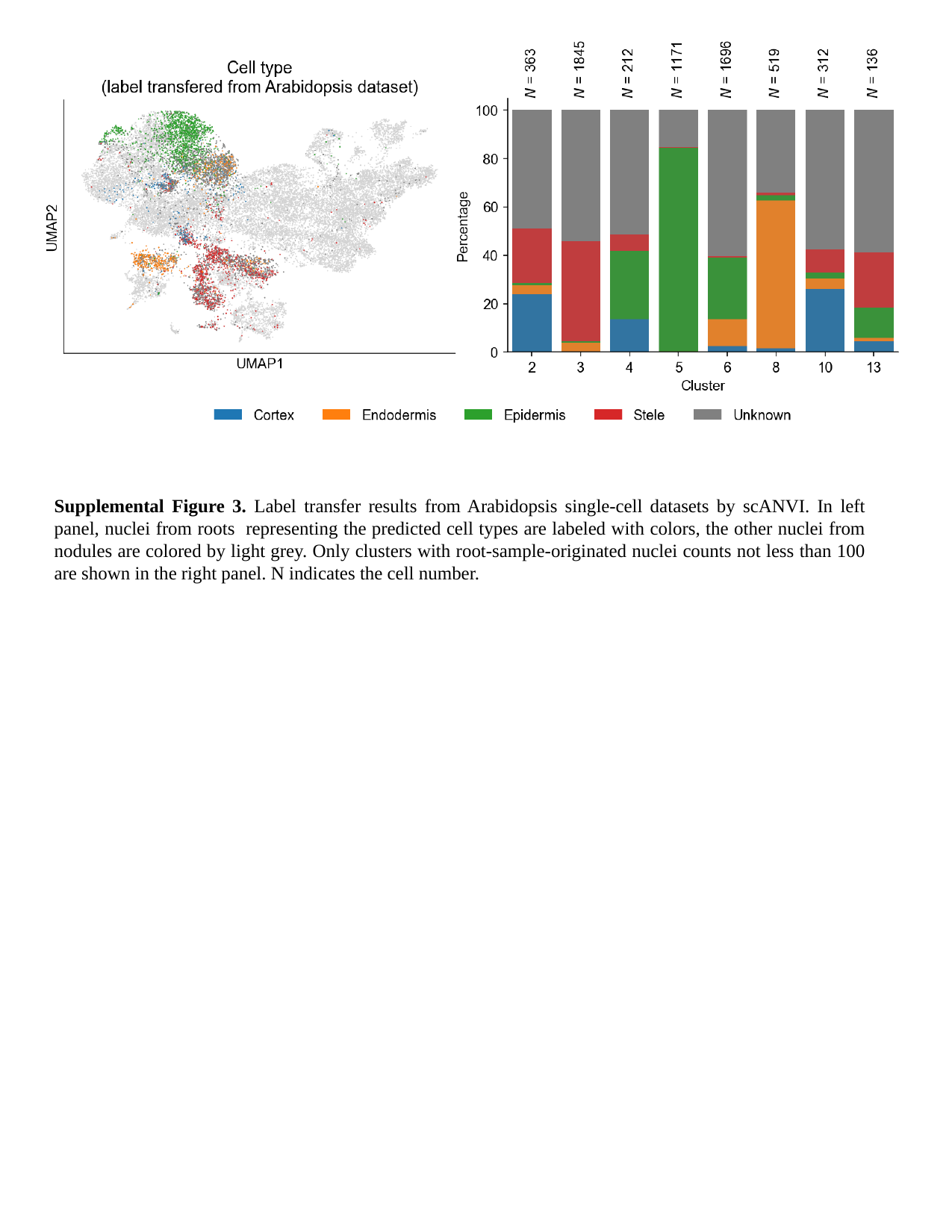

Supplemental Figure 3. Label transfer results from Arabidopsis single-cell datasets by scANVI. In left panel, nuclei from roots representing the predicted cell types are labeled with colors, the other nuclei from nodules are colored by light grey. Only clusters with root-sample-originated nuclei counts not less than 100 are shown in the right panel. N indicates the cell number.

### Slide 4
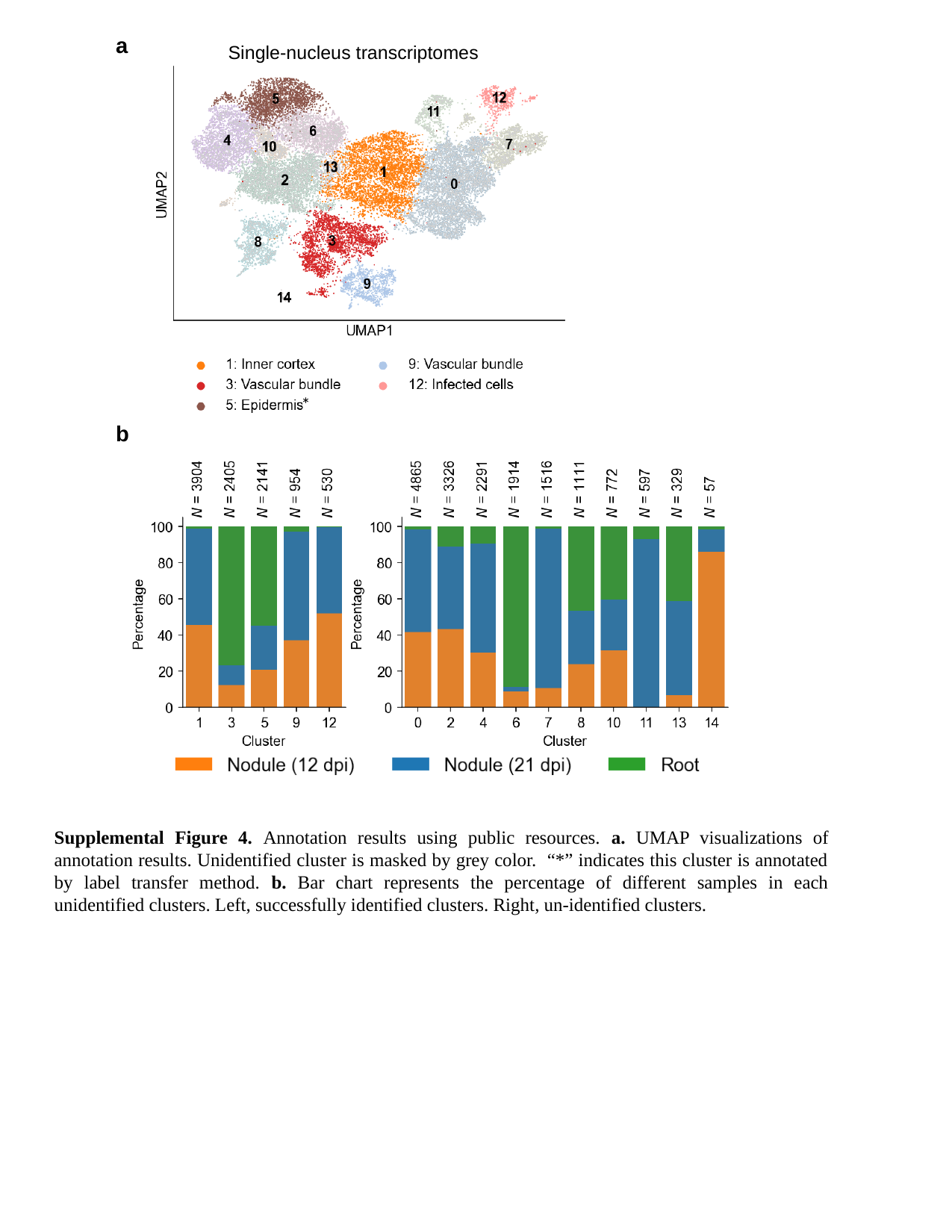

a
Single-nucleus transcriptomes
*
b
Supplemental Figure 4. Annotation results using public resources. a. UMAP visualizations of annotation results. Unidentified cluster is masked by grey color. “*” indicates this cluster is annotated by label transfer method. b. Bar chart represents the percentage of different samples in each unidentified clusters. Left, successfully identified clusters. Right, un-identified clusters.

### Slide 5
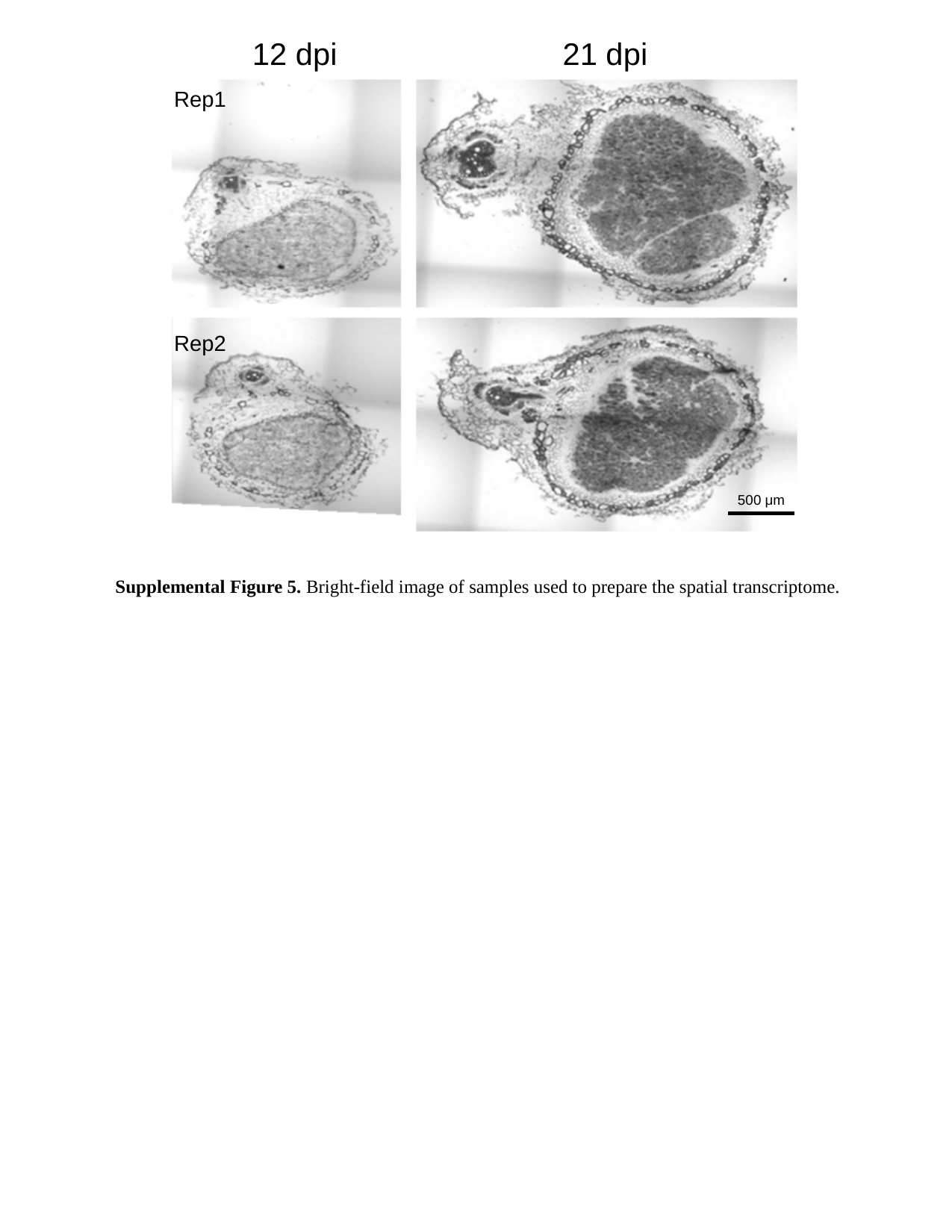

12 dpi
21 dpi
Rep1
Rep2
500 μm
Supplemental Figure 5. Bright-field image of samples used to prepare the spatial transcriptome.

### Slide 6
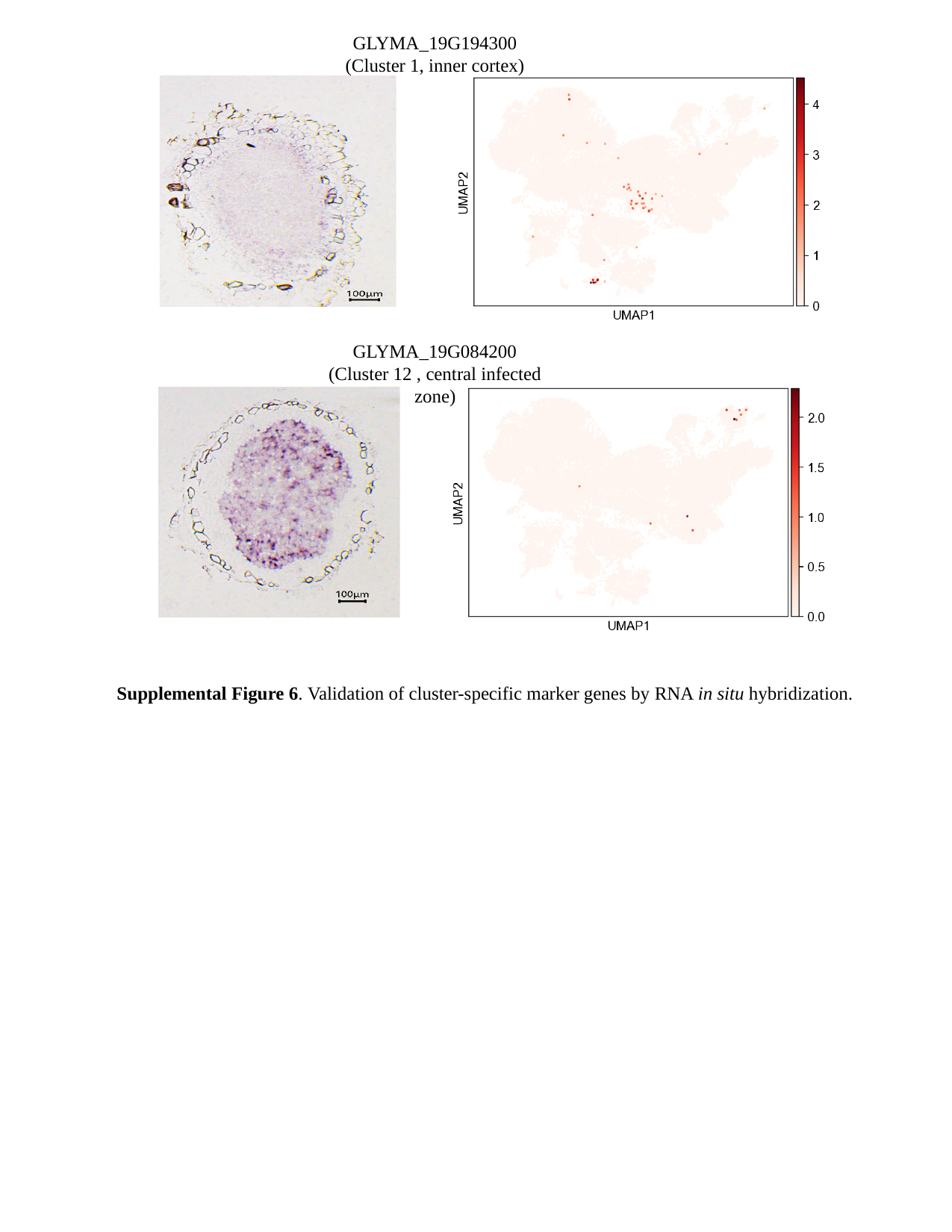

GLYMA_19G194300
(Cluster 1, inner cortex)
GLYMA_19G084200
(Cluster 12 , central infected zone)
Supplemental Figure 6. Validation of cluster-specific marker genes by RNA in situ hybridization.

### Slide 7
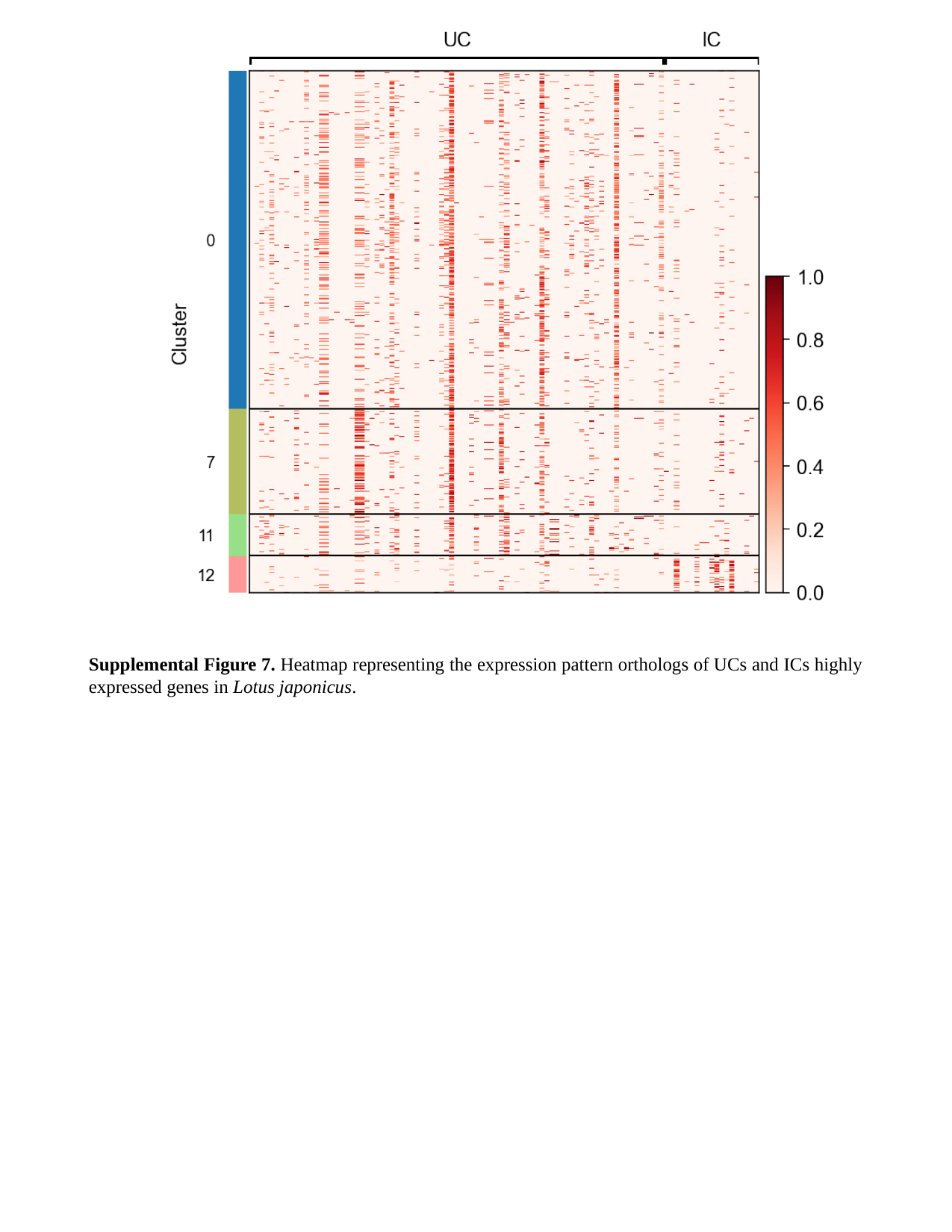

Supplemental Figure 7. Heatmap representing the expression pattern orthologs of UCs and ICs highly expressed genes in Lotus japonicus.

### Slide 8
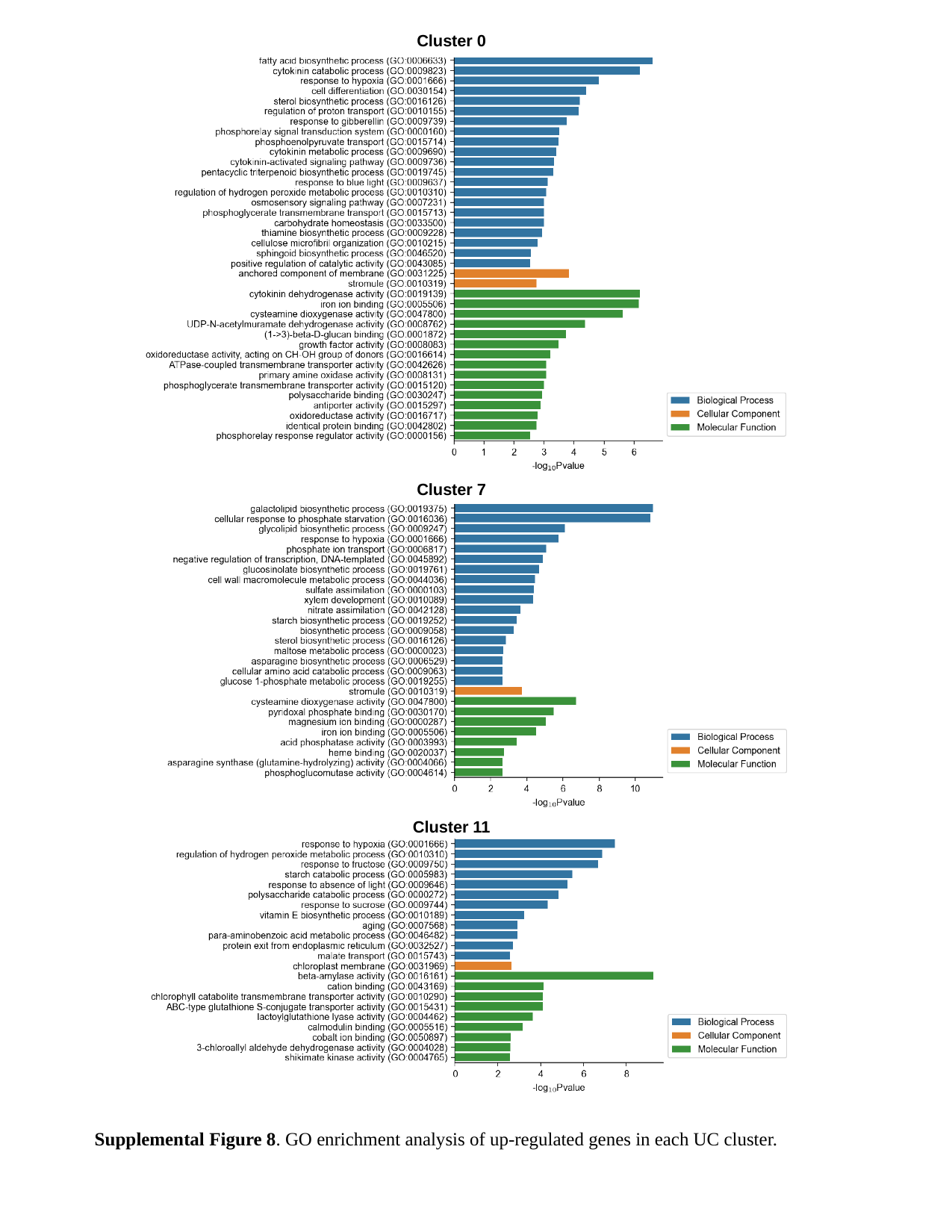

Cluster 0
Cluster 7
Cluster 11
Supplemental Figure 8. GO enrichment analysis of up-regulated genes in each UC cluster.

### Slide 9
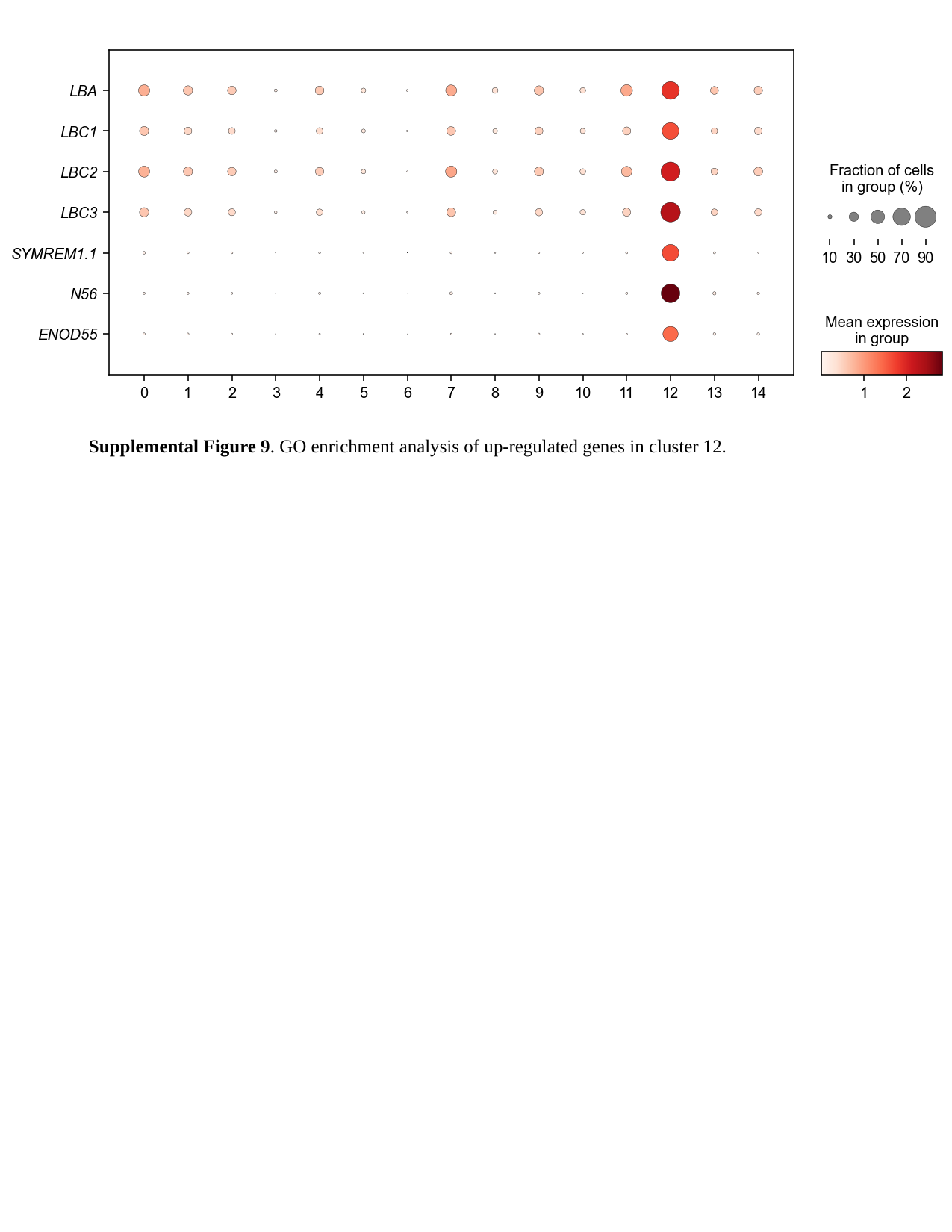

Supplemental Figure 9. GO enrichment analysis of up-regulated genes in cluster 12.

### Slide 10
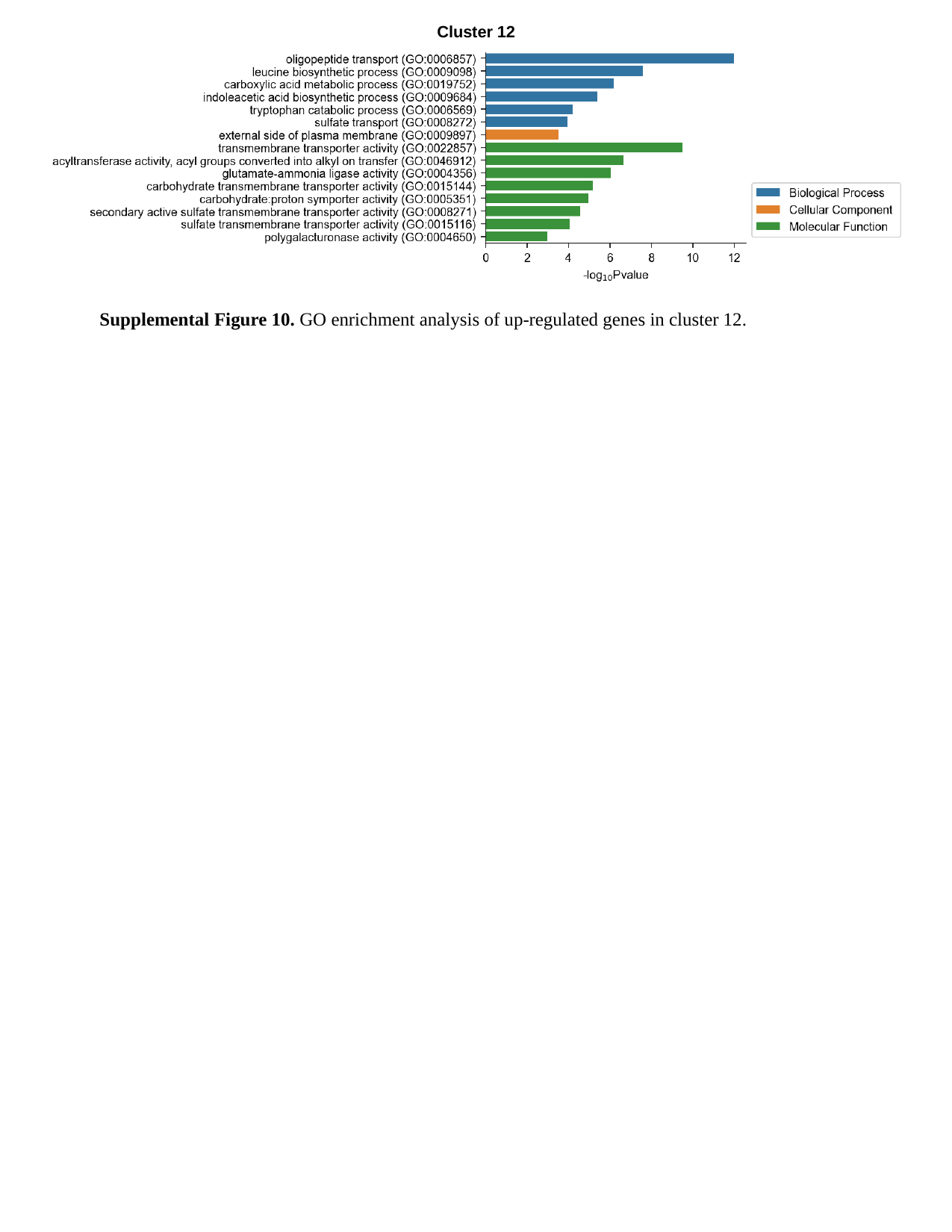

Cluster 12
Supplemental Figure 10. GO enrichment analysis of up-regulated genes in cluster 12.

### Slide 11
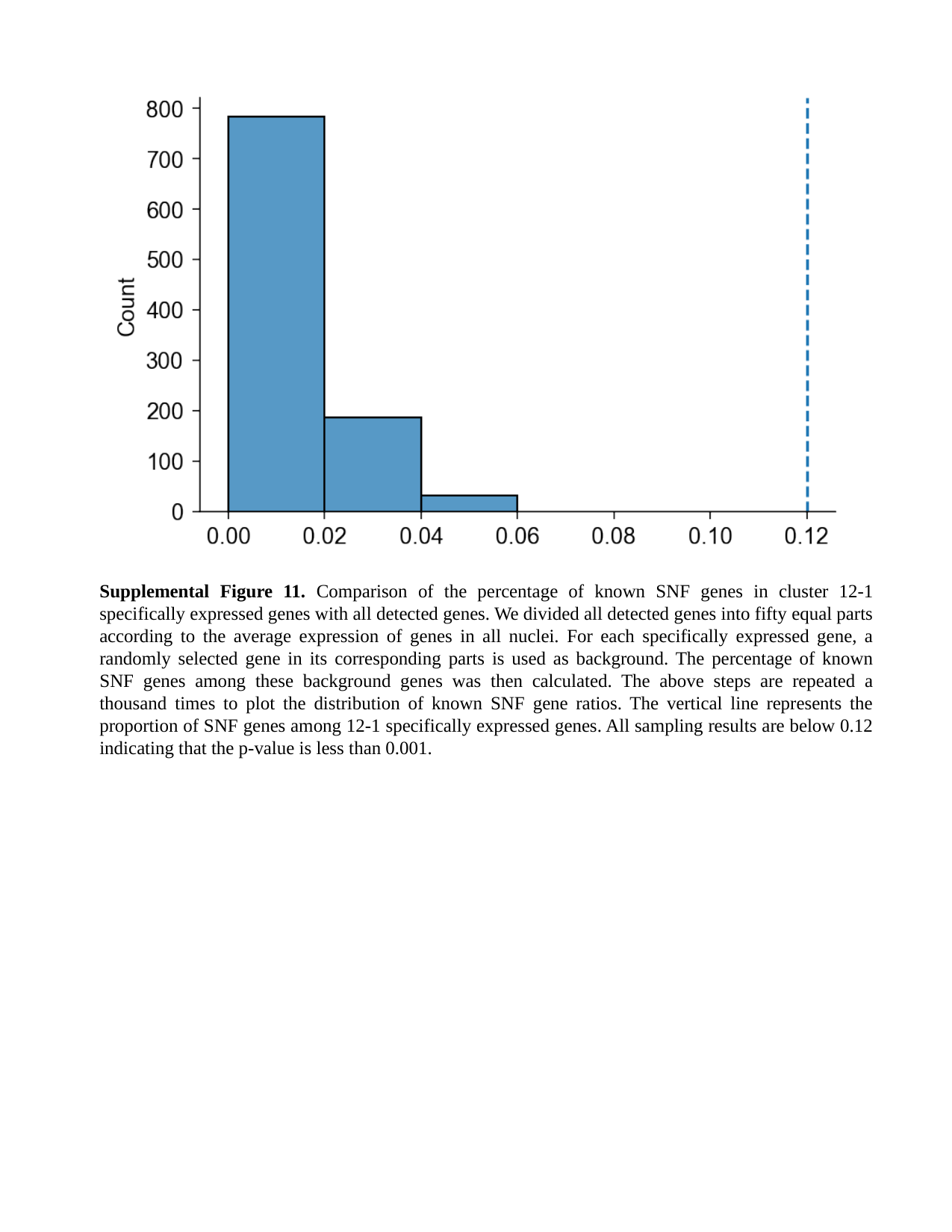

Supplemental Figure 11. Comparison of the percentage of known SNF genes in cluster 12-1 specifically expressed genes with all detected genes. We divided all detected genes into fifty equal parts according to the average expression of genes in all nuclei. For each specifically expressed gene, a randomly selected gene in its corresponding parts is used as background. The percentage of known SNF genes among these background genes was then calculated. The above steps are repeated a thousand times to plot the distribution of known SNF gene ratios. The vertical line represents the proportion of SNF genes among 12-1 specifically expressed genes. All sampling results are below 0.12 indicating that the p-value is less than 0.001.
