## supplemental materials for "Integrated single-nucleus and spatial transcriptomics captures transitional states in soybean nodule symbiosis establishment"

**Materials and Methods**

***Plant growth and nodulation***

The wild-type soybean (Glycine max L. cv Williams 82) seeds were disinfected with chlorine (100 ml NaClO + 4 ml concentrated HCl) and grown on moist sterile filter paper at 22 degree in the dark for 3 days. After germination, the seeds were transferred to pots filled with sterile mixed vermiculite and perlite (2:1, v/v). Nitrogen-free nutrient solution (0.5 mM MgSO_4_,0.2 mM CaCl_2_,0.15 mM K_2_HPO_4_,1 mM K_2_SO_4_,0.02 mM FeCl_3,_ 0.5 µM H_3_BO_3_, 0.1 µM MnSO_4_, 0.15 µM ZnSO_4_, 0.04 µM CuSO_4_, 2.5 pM NaMoO_4_, 2.5 pM CoCl_2_, and 2.5 pM NiSO_4_ included) was poured twice a week. After the cotyledon is spread out, the roots were infected with rhizobium strain USDA110 for nodulation.

***Hairy root transformation***

Agrobacterium rhizogenes-mediated transformation was performed as previously reported^1^. The binary vectors harboring gene construct of interest and a GFP label which can indicate the transgenic roots were introduced into *Agrobacterium rhizogenes* K599. The primary root was cut off at 1 cm below the cotyledons and excised from 7-day-old Soybean (WS82) seedling after the real leaves were unfolded. After the inoculation with *Agrobacterium*, the infected seedlings were placed into the moist sterile vermiculite. The GFP-negative induced hairy roots were removed every seven days, and the transgenic positive roots were retained until the plants had robust transgenic roots that could sustain plant growth (approximately two weeks). Plants were then infected with rhizobia USDA110 for nodulation.

***Histochemical analysis of GUS activity***

~ 2 kb promoter fragments (upstream of ATG) of marker genes identified by single nucleus RNA sequencing were amplified (with primers in Supplemental Data 5) and cloned into the GFP-labelled binary vectors to generate the promoter:GUS constructs. The resulting plasmids were introduced into soybean (WS82) by *Agrobacterium* K599 as described above. The GUS staining were performed as previously described^2,3^. In brief, the fresh nodules were immerged in 5-bromo-4-chloro-3-indolyl-b-D-glucuronic acid (X-Gluc) containing solution and vacuum-infiltrated for 30 min. Histochemical staining for GUS activity was performed at 37 °C for 12 h. After staining, the nodules were fixed with FAA buffer (5% formaldehyde, 5% acetic acid and 50% ethanol) overnight, washed with ethanol and embedded in paraffin. Then, the nodule was sliced transversely into 25-35 µm sections with a microtome (ZEEDO, HS-3315). GUS activity was observed with a light microscope (BSP-8N) equipped with a camera.

***RNA In Situ Hybridization***

Nodules collected at 12 and 21 dpi were fixed in FAA solution for 24 h at 4°C and dehydrated in an ethanol series, cleared in a xylene series, and embedded in paraffin. Then, 10 µm sections were prepared using a microtome (ZEEDO, HS-3315). For probe labeling, specific 500 bp sequences of target genes were amplified from the cDNA of WS82 with primers (Supplemental Data 5) and cloned to pEASY®-Blunt Cloning Vector (TransGen Biotech., #CB111-01). With the resulting vectors as templates, both SP6 and T7 promoter-fused fragments were amplified (with primers in Supplemental Data 5) and purified. Then the digoxigenin-labeled antisense and sense probes were in vitro transcribed with these fragments as templates using SP6 and T7 RNA polymerase with a DIG RNA labeling kit (Roche, 11175025910), respectively. The paraffin on the sections was dissolved with xylene, and the tissue was digested with Proteinase K (Sigma, P2308) and dehydrated with graded alcohol. The prepared probe buffer was added to the tissue sections and incubated for 12 h at 55°C in a humidified box. After hybridization, the sections were incubated with anti-digoxin antibody (Roche, 11093274910) for 90 minutes. Then the sections were washed and incubated in a color reaction solution (Roche, 11681451001) containing blue tetrazolium chloride (NBT) and 5-bromo-4-chloro-3-indolylphosphate (BCIP) for 36 h in a dark humidified box. Photographs were taken under a bright field illumination using a microscope (BSP-8N).

***Nuclei isolation and 10× Single-nucleus RNA-seq library construction***

The 12- and 21-dpi fresh nodules and roots at 21 dpi in the vicinity of mature nodules were collected for single nucleus RNA-seq. The nuclei isolation of root and nodules were performed as previously reported^4^. In brief, the root and nodules were chopped in ice-cold 1× Nuclei isolation buffer (NIB, MilliporeSigma, cat. no. CELLYTPN1) with 1 mM dithiothreitol (DTT, Thermos, R0861), 1× protease inhibitor (Sigma, 4693132001) and 0.4 U/μl Murine RNase inhibitor (Vazyme, R301-03). Then the lysate was filtered with a pre-wet 40 μm strainer and centrifuged at 500 g for 5 min at 4 ℃. The nuclei pellet was resuspended with 500 μl NIB. For sorting, the nuclei were stained with 4,6-Diamidino-2-phenylindole (DAPI) and loaded into a flow cytometer with a 100 μm nozzle. 1 ml 1× PBS with 1% BSA and 0.4 U/μl Murine RNase inhibitor was used as the collection buffer. At least 100,000 nuclei were collected based on the DAPI signal and the nuclear size. The sorted nuclei were pelleted at 4 ℃, 500 g, 5 min, and then resuspended in 50 μl 1× PBS with 1% BSA and 0.4 U/μl Murine RNase inhibitor. After checking the quality of nuclei and counting under a microscope using the DAPI channel, 20000-30000 nuclei were loaded onto the 10x Genomics Chip. The library construction for Illumina sequencing was carried out as described previously^5^.

***Single-nucleus data analysis***

Raw reads were mapped to the Glycine_max_v2.1 reference genome^6^ by Cell Ranger (v6.0.0) using the default parameters except the “include-introns” option is enabled. The abundance matrix was subsequently loaded by SCANPY package^7^ for analysis. For quality control purpose, genes expressed in less than ten nuclei were discarded, only cells with gene counts between 400 and 4000, and UMI counts between 600 and 6000 were kept. Putative doublets were removed by ScDblFinder^8^. The resulted matrixes were then integrated by scVI^9^. We therefore perform leiden algorithm on nearest neighbor graph built on scVI lower-dimension space for clustering and use the UMAP algorithm to visualize the distribution of the data in scVI space. We used existing experimentally validated marker genes (Supplemental Data 3) to unveil the identity of each cell cluster and used the scANVI algorithm^10^ to validate the annotation results using public single-cell data of Arabidopsis roots^11^. One-to-one orthologs were identified by OrthoFinder^12^.

We identified up-regulated genes and specifically expressed genes of each cell cluster using the cellex algorithm^13^. Up-regulated genes are defined as having a specificity score greater than 0.75 and detectable expression in at least 10% of the cells in the corresponding cluster. For cluster-specific expressed gene identified, genes are filtered if they are expressed in less than 20% of cells of the corresponding clusters or more than 1% in the rest of the cells. Then, genes ranked in the top 50 in terms of specificity score were retained.

In trajectory inference step, we combined cellrank^14^ and CytoTRACE^15^ to track the dynamic changes of UCs. The details can be found in their manual (<https://cellrank.readthedocs.io/en/stable>).

In all the above steps, we the AUCell package^16^ to calculate the AUC score of gene set, use clusterProfiler^17^ package to perform GO enrichment analysis and rpy2 package to implement invocation of the R package.

***Stereo-seq and data processing***

Fresh nodules at 12 and 21 dpi were used for Stereo-seq analysis. The Stereo-seq chip preparation and sequencing were performed at the Beijing Genomics Institute (BGI) as previously reported^18^. The raw data was firstly preprocessed by SAW to generated spot-gene matrix^18^. Stained images of sections under bright field are overlayed onto the resulting matrix. Then we performed deconvolution based on single-nucleus sequencing data using destVI^19^. In this step, those clusters with nuclei counts of nodules below 500 were removed.

Reference

1.	Fan, Y.-l. *et al.* One-step generation of composite soybean plants with transgenic roots by Agrobacterium rhizogenes-mediated transformation. *BMC plant biology* **20**, 1-11 (2020).

2.	Zhao, Y., Wang, T., Zhang, W. & Li, X. SOS3 mediates lateral root development under low salt stress through regulation of auxin redistribution and maxima in Arabidopsis. *New Phytologist* **189**, 1122-1134 (2011).

3.	Oh, H.-S. *et al.* The Bradyrhizobium japonicum hsfA gene exhibits a unique developmental expression pattern in cowpea nodules. *Molecular plant-microbe interactions* **14**, 1286-1292 (2001).

4.	Thibivilliers, S., Anderson, D. & Libault, M. Isolation of plant root nuclei for single cell RNA sequencing. *Current protocols in plant biology* **5**, e20120 (2020).

5.	Long, Y. *et al.* FlsnRNA-seq: protoplasting-free full-length single-nucleus RNA profiling in plants. *Genome biology* **22**, 1-14 (2021).

6.	Schmutz, J. *et al.* Genome sequence of the palaeopolyploid soybean. *nature* **463**, 178-183 (2010).

7.	Wolf, F.A., Angerer, P. & Theis, F.J. SCANPY: large-scale single-cell gene expression data analysis. *Genome biology* **19**, 1-5 (2018).

8.	Germain, P.-L., Lun, A., Macnair, W. & Robinson, M.D. Doublet identification in single-cell sequencing data using scDblFinder. *F1000Research* **10**, 979 (2022).

9.	Lopez, R., Regier, J., Cole, M.B., Jordan, M.I. & Yosef, N. Deep generative modeling for single-cell transcriptomics. *Nature methods* **15**, 1053-1058 (2018).

10.	Xu, C. *et al.* Probabilistic harmonization and annotation of single‐cell transcriptomics data with deep generative models. *Molecular systems biology* **17**, e9620 (2021).

11.	Shahan, R. *et al.* A single-cell Arabidopsis root atlas reveals developmental trajectories in wild-type and cell identity mutants. *Developmental cell* **57**, 543-560. e9 (2022).

12.	Emms, D.M. & Kelly, S. OrthoFinder: phylogenetic orthology inference for comparative genomics. *Genome biology* **20**, 1-14 (2019).

13.	Timshel, P.N., Thompson, J.J. & Pers, T.H. Genetic mapping of etiologic brain cell types for obesity. *Elife* **9**(2020).

14.	Lange, M. *et al.* CellRank for directed single-cell fate mapping. *Nature methods* **19**, 159-170 (2022).

15.	Gulati, G.S. *et al.* Single-cell transcriptional diversity is a hallmark of developmental potential. *Science* **367**, 405-411 (2020).

16.	Aibar, S. *et al.* SCENIC: single-cell regulatory network inference and clustering. *Nature methods* **14**, 1083-1086 (2017).

17.	Wu, T. *et al.* clusterProfiler 4.0: A universal enrichment tool for interpreting omics data. *The Innovation* **2**, 100141 (2021).

18.	Xia, K. *et al.* The single-cell stereo-seq reveals region-specific cell subtypes and transcriptome profiling in Arabidopsis leaves. *Developmental Cell* **57**, 1299-1310. e4 (2022).

19.	Lopez, R. *et al.* DestVI identifies continuums of cell types in spatial transcriptomics data. *Nature biotechnology*, 1-10 (2022).
